# Engineering a Decoy Substrate in Soybean to Enable Recognition of the *Soybean Mosaic Virus* NIa Protease

**DOI:** 10.1101/484626

**Authors:** Matthew Helm, Mingsheng Qi, Shayan Sarkar, Haiyue Yu, Steven A. Whitham, Roger W. Innes

**Author notes:** Corresponding author: Roger W. Innes, Department of Biology, Indiana University, Myers Hall 316B, Bloomington, IN 47405, USA.

## Abstract

In Arabidopsis, recognition of the AvrPphB effector protease from *Pseudomonas syringae* is mediated by the disease resistance (R) protein RPS5, which is activated by AvrPphB-induced cleavage of the Arabidopsis protein kinase PBS1. The recognition specificity of RPS5 can be altered by substituting the AvrPphB cleavage site within PBS1 with cleavage sequences for other proteases, including proteases from viruses. AvrPphB also activates defense responses in soybean (*Glycine max*), suggesting that soybean may contain an R protein analogous to RPS5. It was unknown, however, whether this response is mediated by cleavage of a soybean PBS1-like protein. Here we show that soybean contains three *PBS1* orthologs and that their products are cleaved by AvrPphB. Further, transient expression of soybean PBS1 derivatives containing a five-alanine insertion at their AvrPphB cleavage sites activated cell death in soybean protoplasts, demonstrating that soybean likely contains an AvrPphB-specific resistance protein that is activated by a conformational change in soybean PBS1 proteins. Significantly, we show that a soybean PBS1 decoy protein modified to contain a cleavage site for the *Soybean mosaic virus* (SMV) NIa protease triggers cell death in soybean protoplasts when cleaved by this protease, indicating that the PBS1 decoy approach will work in soybean using endogenous *PBS1* genes. Lastly, we show that activation of the AvrPphB-dependent cell death response effectively inhibits systemic spread of SMV in soybean. These data also indicate that decoy engineering may be feasible in other crop plant species that recognize AvrPphB protease activity.

## Introduction

‘Decoy’ engineering is an emerging approach that aims to expand the recognition specificity of intracellular disease resistance proteins in order to generate entirely novel recognition specificities. In this approach, a host protein is engineered to function as a substrate for pathogen-derived effectors (i.e. a decoy) (Harris *et al*., 2013; Segretin *et al*., 2014; Stirnweis *et al*., 2014; Giannakopoulou *et al*., 2015; Maqbool *et al*., 2015). Effector-dependent modification of the decoy triggers activation of an intracellular disease resistance protein, culminating in a hypersensitive response (HR) and disease resistance (Harris *et al*., 2013; Segretin *et al*., 2014; Stirnweis *et al*., 2014; Giannakopoulou *et al*., 2015; Maqbool *et al*., 2015). An example of using decoys to expand the recognition spectrum of an intracellular disease resistance protein is the Arabidopsis RPS5-PBS1 recognition module (Kim *et al*., 2016). In this system, Arabidopsis PBS1 functions as a substrate for the *P. syringae* pv. *phaseolicola* cysteine protease, AvrPphB (Zhu *et al*., 2004). Cleavage of Arabidopsis PBS1 by AvrPphB activates the Arabidopsis <u>n</u>ucleotide-binding leucine-<u>r</u>ich repeat protein (NLR), RPS5, which confers resistance to *P. syringae* (Shao *et al*., 2003; Ade *et al*., 2007; DeYoung *et al*., 2012). Kim *et al*. (2016) demonstrated that the AvrPphB cleavage site sequence within Arabidopsis PBS1 can be substituted with a protease cleavage site sequence recognized by other pathogen-derived proteases, thereby generating a synthetic PBS1 decoy. Protease-dependent cleavage of the PBS1 decoy enables activation of RPS5, which was demonstrated for proteases derived from both bacteria and viruses (Kim *et al*., 2016). These findings thus provide compelling evidence that engineering decoys based on the Arabidopsis RPS5-PBS1 recognition module may be an effective NLR gene-based strategy to control plant diseases in crop plants.

Creation of a decoy recognition system in crop plants based on PBS1 may not require use of Arabidopsis genes. Arabidopsis *PBS1* is a well-conserved defense gene, with orthologs present in monocot and dicot crop plant species (Caldwell and Michelmore, 2009). Importantly, AvrPphB has been shown to cleave PBS1 orthologs from both wheat and barley, and to induce an HR in these species, as well as in soybean (Russell *et al*., 2015; Sun *et al*., 2017; Carter *et al*., in press). Carter *et al*. (in press) recently mapped the AvrPphB response in barley to a single locus containing an NLR gene, *Avr<u>P</u>ph<u>B</u> <u>R</u>esistance <u>1</u>* (*Pbr1*). Significantly, PBR1 co-immunoprecipitates with barley and *N. benthamiana* PBS1 proteins and co-expression of PBR1 with AvrPphB activates a cell death response in *N. benthamiana* (Carter *et al*., in press). It is thus likely that other crop plants that recognize AvrPphB protease activity also contain an AvrPphB-specific resistance protein that guards PBS1 orthologous proteins.

In the present study, we sought to generate PBS1-based decoys in soybean that would confer recognition of the NIa protease from *Soybean mosaic virus* (SMV; genus *Potyvirus*). SMV is the most widespread virus that infects soybean and is responsible for significant economic losses worldwide (Whitham *et al*., 2016; Hajimorad *et al*., 2018). In addition, the prevalence and severity of losses to SMV in the United States have increased over the last two decades, which has been primarily attributed to the introduction of the soybean-colonizing aphid (*Aphis glycines*), a vector for SMV (Hartman *et al*., 2001; Hill *et al*., 2001; Clark and Perry, 2002). SMV is a single-stranded, positive-sense filamentous RNA virus (Whitham *et al*., 2016; Hajimorad *et al*., 2018). Upon SMV infection, the viral RNA is translated as a precursor polyprotein that is proteolytically processed by three SMV-encoded proteases at internal cleavage sites to produce mature, multifunctional viral proteins, including P1 (protein 1), HC-Pro (helper component protease), P3 (protein 3), 6K1 (six kiloDalton 1), CI (cylindrical inclusion), 6K2 (six kiloDalton 2), NIa (nuclear inclusion a), NIb (nuclear inclusion b), and CP (coat protein) (Hajimorad *et al*., 2018). Significantly, the NIa protease is the only SMV-encoded protease that acts in *trans* (Adams *et al*., 2005). Further, the minimal amino acid sequence required for recognition by the SMV NIa protease has been previously characterized and is well conserved among SMV isolates (Ghabrial *et al*., 1990; Jayaram *et al*., 1992; Adams *et al*., 2005). *Potyvirus* proteases are essential for processing the viral polyprotein into functional viral proteins (Adams *et al*., 2005). We, therefore, hypothesize that a resistance protein activated by the enzymatic activity of the NIa protease would be a durable disease resistance trait as it would be unlikely SMV would simultaneously change specificity of the NIa protease and multiple protease cleavage sites embedded within the polyprotein. The observation that soybean responds to AvrPphB with a hypersensitive response (Russell *et al*., 2015) suggests that artificial soybean PBS1-based decoys can be engineered to detect the NIa protease from SMV. It was unclear, however, whether the endogenous soybean resistance protein that detects AvrPphB protease activity functions analogously to Arabidopsis RPS5.

Here, we show that soybean contains three plasma membrane-localized PBS1 orthologous proteins (*Gm*PBS1-1, *Gm*PBS1-2, and *Gm*PBS1-3) that are cleaved by AvrPphB. Significantly, transient expression of *Gm*PBS1 derivatives containing a five alanine insertion at the AvrPphB cleavage site (*Gm*PBS1^5Ala^) induces cell death in the absence of AvrPphB, demonstrating that *Gm*PBS1 proteins have a functional role in the innate immune response, likely by being guarded by an NLR protein functionally analogous to RPS5. Significantly, we demonstrate that replacing the native AvrPphB cleavage site sequence with a SMV NIa protease recognition site in *Gm*PBS1-1 (*Gm*PBS1-1^SMV^) results in NIa-mediated cleavage, and such cleavage activates cell death in soybean protoplasts. Lastly, we show that SMV-mediated overexpression of AvrPphB inhibits systemic spread of SMV in soybean, demonstrating that the AvrPphB-dependent cell death response resulting from *Gm*PBS1 cleavage is effective against a viral pathogen. Collectively, these data suggest that synthetic PBS1-based decoys can be used to expand effector protease recognition in soybean and generating artificial decoys offers an attractive approach for engineering resistance to other soybean pathogens.

## Results

### Soybean contains three *PBS1* genes that are co-orthologous to *Arabidopsis PBS1* and whose protein products are cleaved by AvrPphB

*Pseudomonas syringae* pv. *glycinea* Race 4 (PsgR4) expressing the effector protease AvrPphB elicits a hypersensitive response in soybean (*Glycine max*), indicating that soybean contains an AvrPphB-specific disease resistance protein (Russell *et al*., 2015). To confirm these observations, we delivered AvrPphB or an enzymatically inactive derivative of AvrPphB, AvrPphB(C98S), to primary leaves of soybean (cv. Flambeau) using *P. syringae* pathovar *tomato* strain D36E, which lacks all known endogenous type III effectors (Wei *et al*., 2015; Carter *et al*., in press). Consistent with the observations of Russell *et al*. (2015), D36E(AvrPphB) induced an observable cell death response 24 hours post-injection (hpi), while minimal cell death was observed with D36E(C98S), or D36E carrying the empty vector (Supp. Fig. S1). These data indicate that soybean likely contains a disease resistance protein that can detect the protease activity of AvrPphB.

Given that Arabidopsis detects AvrPphB protease activity via sensing cleavage of the protein kinase PBS1, we hypothesized that soybean may employ a similar mechanism. We thus screened for soybean PBS1 homologs that can be cleaved by AvrPphB. Using the Arabidopsis PBS1 amino acid sequence (*At*PBS1) as a query, we used BLAST to identify the top twenty soybean PBS1-like (*Gm*PBL) protein sequences (release Williams82.a2.v1; http://soybase.org) (Grant *et al*., 2010) with the most similarity to *At*PBS1. Phylogenetic analysis showed that Glyma.08G360600, Glyma.10G298400, and Glyma.20G249600 are more closely related to *At*PBS1 than to other Arabidopsis or soybean PBL proteins (Fig. 1A; Supp. Fig. S2). Full-length amino acid alignments showed that Glyma.08G360600, Glyma.10G298400, and Glyma.20G249600 are 80%, 77%, and 77% identical to *At*PBS1, respectively (Supp. Fig. 3), with alignment across just the kinase domains showing even higher identities (91%, 92%, and 92%). Based on the structure of the phylogenetic tree, all three soybean genes are co-orthologous to *AtPBS1*. We therefore designated Glyma.08G360600 as *Gm*PBS1-1 (GenBank: MK035866), Glyma.10G298400 as *Gm*PBS1-2 (GenBank: MK035867), and Glyma.20G249600 as *Gm*PBS1-3 (GenBank: MK035868).

**Figure 1.**
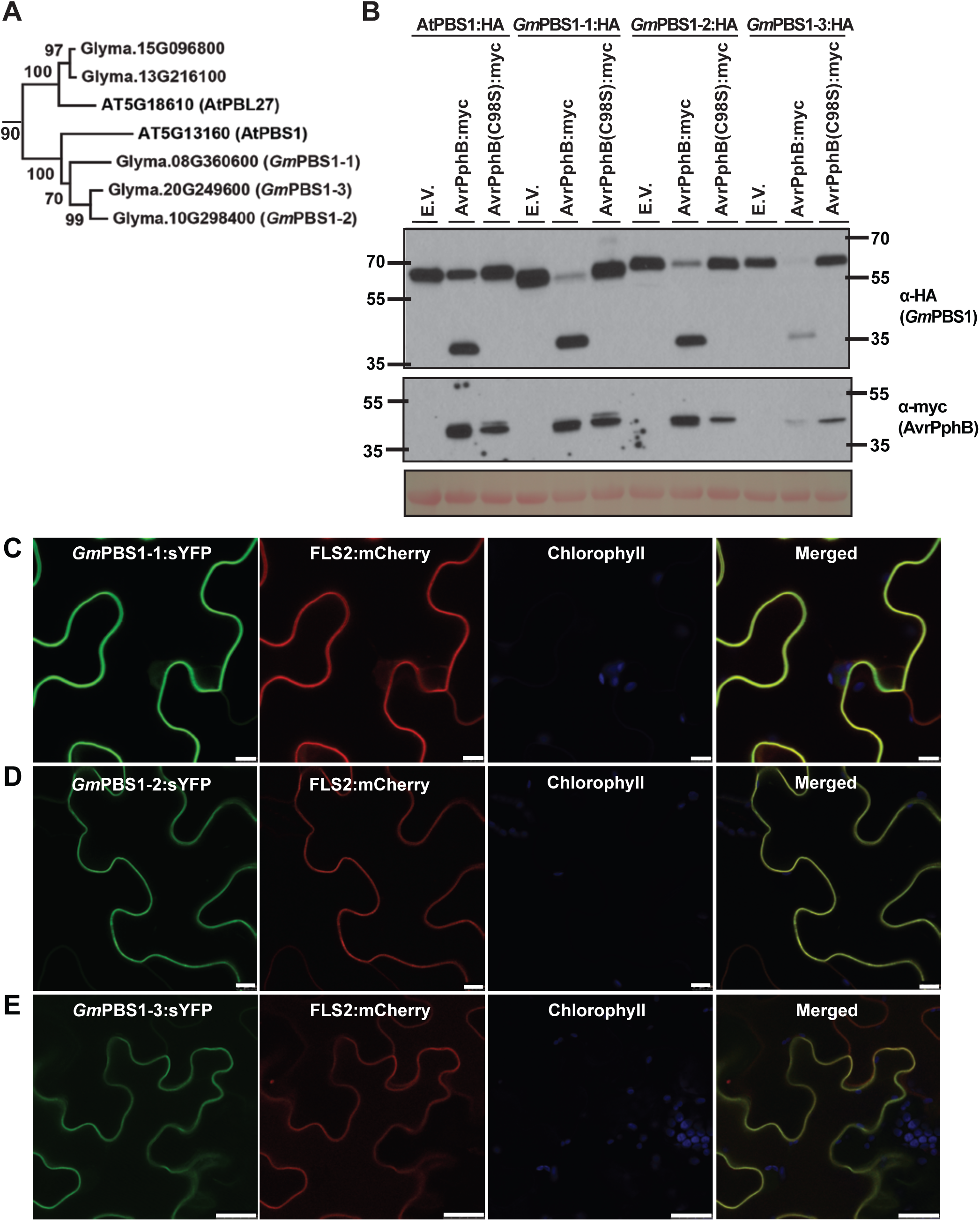
Soybean contains three PBS1 proteins that localize to the plasma membrane and are cleaved by AvrPphB. A) Glyma.08G360600.3 (*Gm*PBS1-1), Glyma.10G298400.1 (*Gm*PBS1-2), and Glyma.20G249600.2 (*Gm*PBS1-3) are co-orthologous to Arabidopsis PBS1 (*At*PBS1). Shown is a phylogenetic tree generated from the amino acid sequences of Arabidopsis PBS1 and the most closely related soybean homologs using MEGA7 with the neighbor joining model (Kumar et al., 2016). The bootstrap values are shown at the nodes. This tree is a subset of Supplemental Figure 2, which displays soybean proteins closely related to Arabidopsis PBS1 and Arabidopsis PBS1-like (*At*PBL) proteins. B) Cleavage of *Gm*PBS1-1, *Gm*PBS1-2, and *Gm*PBS1-3 by AvrPphB. HA-tagged soybean PBS1 homologs or Arabidopsis PBS1 were transiently co-expressed with or without myc-tagged AvrPphB and AvrPphB(C98S) in *N. benthamiana*. Total protein was extracted six hours post-transgene induction and immunoblotted with the indicated antibodies. Ponceau S solution staining was included as a control to show equal loading of protein samples. Three independent experiments were performed with similar results. The results of only one experiment are shown. (C-E) The soybean PBS1 proteins localize to the plasma membrane (PM) in *N. benthamiana*. C) sYFP-tagged Glyma.08G360600.3 (*Gm*PBS1-1), D) Glyma.10G298400.1 (*Gm*PBS1-2), and E) Glyma.20G249600.2 (*Gm*PBS1-3) and mCherry-tagged FLS2 were transiently co-expressed in *N. benthamiana* leaves. Live-cell imaging was performed using laser-scanning confocal microscopy 24 hours following transgene induction. FLS2 was included as a reference for plasma membrane localization. Scale bars = 10 μm, except in C, in which the bar = 25 μm. Two independent experiments were performed with similar results. The results of only one experiment are shown.

The AvrPphB cleavage site sequence is conserved in all three *Gm*PBS1 proteins (Supp. Fig. 3), suggesting that these proteins should be cleavable by AvrPphB. To test this, *Gm*PBS1-1, *Gm*PBS1-2, and *Gm*PBS1-3 were transiently co-expressed with AvrPphB in *Nicotiana benthamiana*. Immunoblot analysis showed that all three proteins are cleaved by AvrPphB and not by AvrPphB(C98S) (Fig. 1B), indicating that recognition of AvrPphB in soybean could be mediated by cleavage of one, or more, of these three *Gm*PBS1 proteins.

In Arabidopsis, detection of PBS1 cleavage occurs at the plasma membrane, and *At*PBS1 is targeted to the plasma membrane via N-terminal myristoylation and palmitoylation motifs (Qi *et al*., 2014). These motifs are conserved in all three *Gm*PBS1 proteins (Supp. Fig. 3), so we assessed whether these proteins are also targeted to the plasma membrane using transient expression of superYFP-tagged versions in *N. benthamiana.* All three proteins displayed a clear plasma membrane localization pattern, co-localizing with the known plasma membrane protein, *At*FLS2 (Fig. 1C).

### Insertion of five alanine residues in the AvrPphB cleavage site of soybean PBS1 proteins activates cell death in the absence of AvrPphB-mediated cleavage

The above data are consistent with AvrPphB being recognized via cleavage of one or more *Gm*PBS1 proteins, but do not prove it. In Arabidopsis, AvrPphB targets at least nine Arabidopsis PBS1-like (*At*PBL) proteins (Zhang *et al*., 2010; DeYoung *et al*., 2012). It is therefore a formal possibility that soybean detects AvrPphB protease activity by sensing cleavage of an AvrPphB substrate other than *Gm*PBS1 proteins. To assess whether *Gm*PBS1 cleavage does indeed activate resistance in soybean, we inserted five alanine residues at the AvrPphB cleavage site of *Gm*PBS1-1 (*Gm*PBS1-1^5Ala^; Fig. 2A). An equivalent insertion in *At*PBS1 induces a conformational change that activates RPS5-dependent cell death in Arabidopsis in the absence of AvrPphB expression (DeYoung *et al*., 2012). We thus hypothesized that a five-alanine insertion into one of the *Gm*PBS1 proteins would activate the AvrPphB-specific R protein in soybean, and thus induce cell death. We selected *Gm*PBS1-1 for this assay because it is the most abundantly expressed of the three *GmPBS1* co-orthologs in leaves (Libault *et al*., 2010). We then transiently transfected soybean (cv. Williams 82) protoplasts with either *Gm*PBS1-1 or the *Gm*PBS1-1^5Ala^ derivative along with a *Renilla* luciferase reporter (Fig. 2B). In this assay, a reduction in luciferase activity indicates activation of cell death. As positive controls for cell death, we transiently expressed AvrB or AvrPphB, which activate a hypersensitive response in Williams 82 (Ashfield *et al*., 2004). Consistent with our hypothesis, transient expression of *Gm*PBS1-1^5Ala^, but not wild-type *Gm*PBS1-1, induced cell death similar to that observed with AvrB and AvrPphB, demonstrating that insertion of five alanine residues in the activation loop of *Gm*PBS1-1 activates a cell death response in soybean (Fig. 2B). To test whether the cell death response is specific to *Gm*PBS1-1, we transiently transfected soybean protoplasts with either *Gm*PBS1-2 or *Gm*PBS1-2^5Ala^ and quantified luciferase activity. Transient expression of *Gm*PBS1-2^5Ala^, but not *Gm*PBS1-2, also induced cell death equivalent to AvrB and AvrPphB (Supp. Fig. S4A). Collectively, these data suggest that soybean likely senses AvrPphB protease activity via cleavage of a *Gm*PBS1 protein, analogous to the Arabidopsis RPS5-PBS1 recognition system.

**Figure 2.**
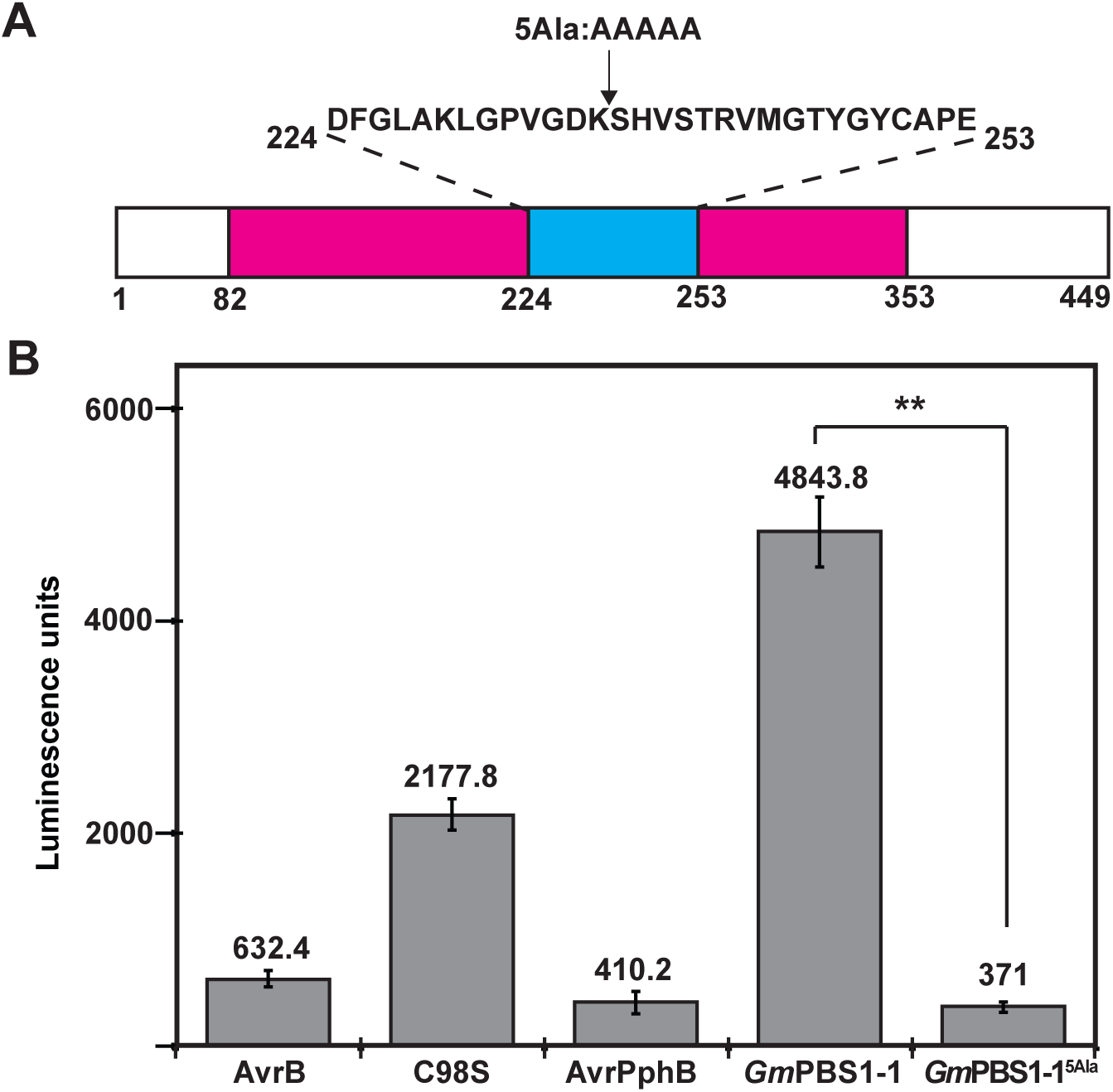
Transient expression of the *Gm*PBS1-1^5Ala^ derivative activates cell death in soybean (cv. Williams 82) A) Schematic illustration of the synthetic *Gm*PBS1-1^5Ala^ construct. The predicted kinase domain (amino acids 82-353) and activation segment (amino acids 224-253) of *Gm*PBS1-1 are represented by a magenta box and a cyan blue box, respectively. The amino acid sequence of the activation segment and the location of the five-alanine insertion are indicated above. B) Transient expression of the *Gm*PBS1-1^5Ala^ derivative activates cell death in soybean protoplasts. The indicated constructs were transiently co-expressed along with *Renilla* Luciferase in soybean (cv. Williams 82) protoplasts. Values represent the mean ± S.D. for two technical replicates. T-tests were performed for the pair-wise comparison. The double asterisk indicates significant difference (p < 0.01). Three independent experiments were performed with similar results. The results of one experiment are shown.

### *Soybean mosaic virus* (SMV) NIa protease-mediated cleavage of *Gm*PBS1-1^SMV^ decoy protein activates cell death in soybean protoplasts

Our evidence suggesting that soybean contains an AvrPphB recognition system functionally analogous to the Arabidopsis RPS5-PBS1 pathway raises the possibility that soybean PBS1 proteins can be modified to enable cleavage by other pathogen proteases, and thus expand the recognition specificity of the AvrPphB-specific R protein in soybean. We have previously shown that *At*PBS1 can be modified to be cleaved by the NIa protease from *Turnip mosaic virus* (TuMV), with transgenic Arabidopsis plants expressing this ‘decoy’ protein displaying enhanced resistance to TuMV (Kim *et al.*, 2016). To create a suitable soybean PBS1 decoy protein for detection of *Soybean mosaic virus* (SMV), we replaced the AvrPphB cleavage site sequence in the activation loop of *Gm*PBS1-1 with a known SMV NIa protease cleavage sequence [ESVLSQS; (Ghabrial *et al*., 1990)] (Fig. 3A). As shown in Fig. 3B, *Gm*PBS1-1^SMV^ is cleaved by SMV NIa protease when transiently co-expressed in *N. benthamiana*, but not by AvrPphB, while wild-type *Gm*PBS1-1 is cleaved by AvrPphB, but not by the SMV NIa protease. We then tested for activation of cell death in soybean cells using the protoplast transformation system described above. Co-expression of the NIa protease with *Gm*PBS1-1^SMV^ resulted in a significant reduction in luciferase activity compared to co-expression with wild-type *Gm*PBS1-1, indicating that NIa-mediated cleavage of the *Gm*PBS1-1^SMV^ decoy activates cell death in soybean cells (Fig. 3C). To test whether *Gm*PBS1-2 and *Gm*PBS1-3 could also serve as decoys, we replaced the AvrPphB cleavage site sequence with the NIa protease cleavage sequence (Supp. Figs. 5A and 5B). Transient co-expression of either the *Gm*PBS1-2^SMV^ or *Gm*PBS1-3^SMV^ with the NIa protease in *N. benthamiana* resulted in NIa-mediated cleavage of the decoy proteins (Supp. Figs. 5C and 5D). These data thus suggest synthetic soybean PBS1 proteins can serve as decoys for the NIa protease from SMV, thereby expanding the recognition specificity of the AvrPphB-specific R protein in soybean.

**Figure 3.**
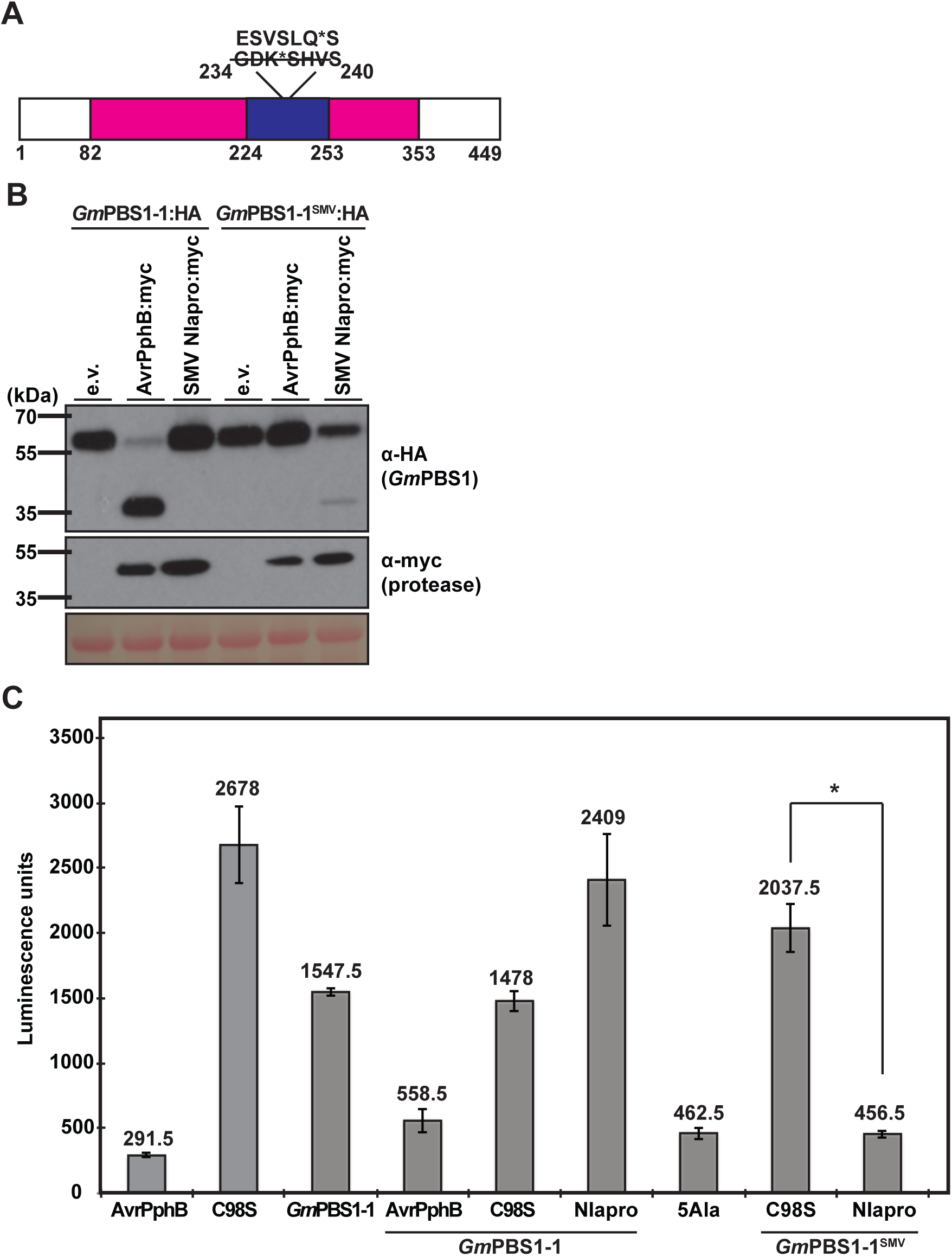
SMV NIa-mediated cleavage of the *Gm*PBS1-1^SMV^ decoy activates cell death in soybean cv. Williams 82. A) Schematic representation of the synthetic *Gm*PBS1-1^SMV^ decoy. The endogenous AvrPphB cleavage site in *Gm*PBS1-1 (GDKSHVS) was substituted with the cleavage site sequence recognized by the SMV NIa protease (ESVSLQS). The asterisks indicate the location of cleavage by the respective proteases within the recognition sites. B) Cleavage of the *Gm*PBS1-1^SMV^ synthetic decoy protein by the SMV NIa protease. HA-tagged *Gm*PBS1-1^SMV^ or *Gm*PBS1-1 were transiently co-expressed with either empty vector (e.v.), AvrPphB:myc, or SMV NIapro:myc in *N. benthamiana*. Total protein was extracted nine hours post-transgene induction and immunoblotted with the indicated antibodies. Ponceau S solution staining was included as a control to show equal loading of protein samples. Three independent experiments were performed with similar results. The results of only one experiment are shown. C) Cleavage of the *Gm*PBS1-1^SMV^ decoy by the NIa protease activates cell death in soybean protoplasts. The indicated constructs were transiently co-expressed along with *Renilla* Luciferase in soybean (cv. Williams 82) protoplasts. Values represent the mean ± S.D. for two technical replicates. T-tests were performed for the pair-wise comparison. The asterisk indicates significant difference (p < 0.05). Two independent experiments were performed with similar results. The results of one experiment are shown.

### Recognition of AvrPphB protease activity in soybean inhibits systemic spread of SMV

Our evidence demonstrating that soybean PBS1 proteins can be engineered to confer recognition of the NIa protease from SMV suggests decoy engineering can be extended into soybean. It is unclear, however, whether the cell death response elicited by AvrPphB protease activity is effective against SMV in soybean. Kim *et al*. (2016) showed that Arabidopsis RPS5 can be activated by sensing cleavage of an engineered PBS1 decoy by the NIa protease from TuMV, thereby broadening its recognition specificity. However, infection of transgenic Arabidopsis expressing the PBS1 decoy protein by TuMV resulted in a lethal systemic necrosis phenotype, demonstrating that RPS5-mediated defense responses confers only partial resistance against TuMV (Kim *et al*., 2016). To test whether activation of the AvrPphB-dependent cell death response could inhibit systemic spread of SMV in soybean, we used an SMV-mediated transient expression system to transiently express green fluorescent protein (GFP), AvrPphB or AvrPphB(C98S) in soybean. Using this approach, (Wang *et al*., 2006) showed that AvrB, an effector from *P. syringae* pv. *glycinea*, activates defense responses and inhibits systemic spread of SMV into the upper, uninoculated trifoliate leaflets of soybean (cv. Harosoy). We inserted the open reading frames (ORFs) encoding AvrPphB and AvrPphB(C98S) into pSMV-Nv (Fig. 4A). Primary leaves of soybean were mechanically inoculated with DNA of either pSMV-Nv::*GFP*, pSMV-Nv::*AvrPphB,* or pSMV-Nv::*AvrPphB*. Consistent with the observations of Wang *et al*. (2006), insertion of the GFP ORF resulted in development of mosaic symptoms and leaf rugosity in the systemic, uninoculated trifoliate leaflets indicative of successful SMV infection (Fig. 4B). In addition to the observed SMV symptoms, immunoblot analysis showed detectable SMV coat protein (SMV CP) and GFP protein accumulation in the systemic, uninoculated fourth trifoliate leaflet (Fig. 4C), demonstrating the recombinant virus did not spontaneously delete the 0.7kb insert and that GFP is stably expressed *in planta*. In contrast, inoculation of leaves with pSMV-Nv::*AvrPphB* did not result in any systemic SMV symptoms, and no AvrPphB or SMV CP accumulation was detected in the fourth trifoliate leaflet three weeks post-inoculation (Figs. 4B and 4C). Expression of pSMV-Nv::*AvrPphB(C98S)*, however, resulted in mosaic symptoms and leaf rugosity similar to that observed with pSMV-Nv::*GFP*, as well as detectable AvrPphB protein accumulation in the fourth trifoliate leaflet (Figs. 4B and 4C). Collectively, these data suggest that activation of the AvrPphB-dependent cell death response effectively inhibits systemic spread of SMV in soybean and, therefore, synthetic decoy engineering may be an effective strategy for engineering resistance to SMV.

**Figure 4.**
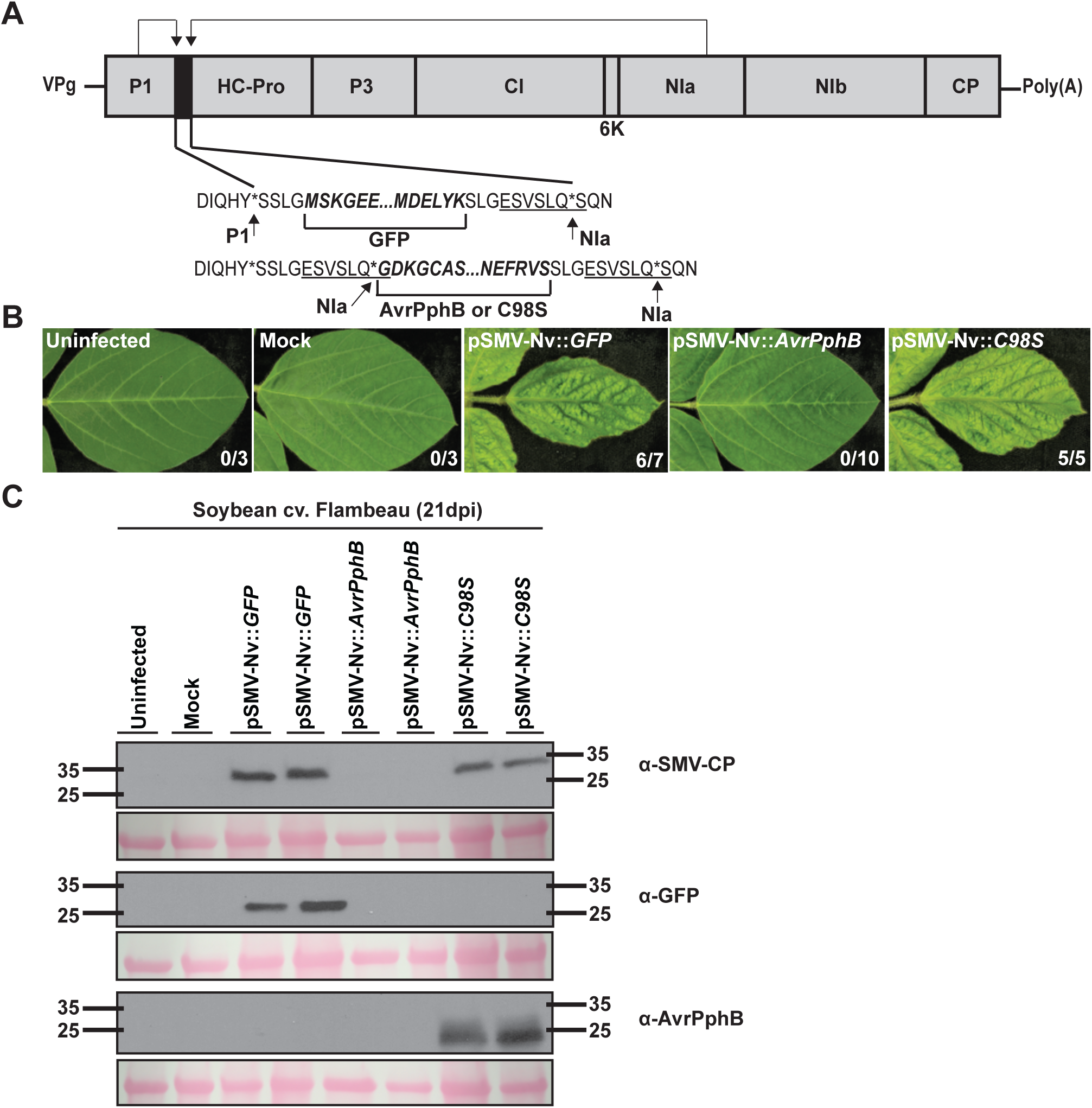
Recognition of AvrPphB protease activity in soybean blocks SMV symptom development and viral protein accumulation in systemic, uninoculated trifoliate leaflets. A) Schematic representation of the SMV-based transient expression system used in this study (adapted from Wang et al., 2006). The grey boxes represent SMV-N cistrons. The shaded black box indicates the location of transgene insertion. Arrows indicate the positions of the P1 and NIa protease cleavage sites within the SMV-Nv polyprotein. Cleavage by the P1 and the NIa proteases at the respective cleavage sites (indicated by the arrows) releases GFP, AvrPphB, or AvrPphB(C98S) from the SMV polyprotein. The SMV NIa protease recognition site is underlined. B) Recognition of AvrPphB protease activity in soybean (cv. Flambeau) inhibits SMV movement into uninoculated trifoliate leaflets. Fourteen-day-old soybean (cv. Flambeau) primary leaves were rub-inoculated with either mock (buffer) or 35S-driven infectious cDNAs of strain SMV-Nv expressing GFP (pSMV-Nv::*GFP*), AvrPphB (pSMV-Nv::*AvrPphB*), or AvrPphB(C98S) (pSMV-Nv::*C98S*). Three weeks post-inoculation, the fourth trifoliate leaflet was photographed under white light. The numbers on the right bottom of the photographs indicate the sum of trifoliate leaflets displaying viral symptoms consistent with SMV infection/total number of plants rub-inoculated with infectious cDNAs. Two independent experiments were performed with similar results. The results of only one experiment are shown. C) Western blot analysis shows SMV coat protein (SMV-CP) accumulation in the systemic trifoliate leaflets of soybean (cv. Flambeau) inoculated with pSMV-Nv::*GFP* and pSMV-Nv::*C98S* and not pSMV-Nv::*AvrPphB*. Three weeks post-inoculation, the fourth trifoliate leaflet was flash frozen in liquid nitrogen, total protein extracted, and protein concentration estimated by Bradford (1976) assay. Ten micrograms of total protein was separated on 4-20% gradient PreciseTM Protein Gels and immunoblotted with the indicated antibodies. Lanes with duplicate labels indicate independent biological replicates. Ponceau S solution staining was included as a control to show equal loading of protein samples. Two independent experiments were performed with similar results. The results of one experiment are shown.

## Discussion

We have previously shown that AvrPphB activates a hypersensitive response in most soybean varieties (Russell *et al*., 2015), but it was unclear whether this response was mediated by cleavage of a PBS1-like protein, and hence whether it would be feasible to use a PBS1 decoy strategy to engineer novel recognition specificities in soybean. To address these questions, we first identified soybean *PBS1* orthologs, and assessed whether the encoded proteins were cleaved by AvrPphB (Fig. 1). These analyses confirmed that *Gm*PBS1 proteins are cleaved by AvrPphB, suggesting that AvrPphB protease activity may be activating NLR-triggered immunity in soybean via a mechanism similar to that employed by Arabidopsis (Ade *et al*., 2007).

To confirm that *Gm*PBS1 modification activates cell death in soybean, we developed a protoplast transformation assay. Research in soybean is often hampered by the lack of rapid, reproducible transient gene expression methods. Our demonstration of reproducible protoplast assays for cell death following *Gm*PBS1 cleavage thus opens up many possibilities for investigating soybean immune signaling. Although routinely used to assess gene function in other plant species, protoplast transformation is often technically challenging, and a robust method for preparation and transformation of soybean protoplasts has only recently been reported (Wu and Hanzawa, 2018). Wu and Hanzawa (2018) demonstrated expression of GFP, and the nuclear localization of the E1 protein (Glyma.06G207800) fused to GFP in soybean protoplasts isolated from the Williams 82 cultivar. Prior to this work, there have been few publications regarding the preparation or use of protoplasts from soybean (Wu and Hanzawa, 2018). These include recent papers by (Sun *et al*., 2015) and (Kim *et al*., 2017) who reported gene editing in protoplasts of the Williams 82 soybean cultivar following delivery of DNA constructs expressing Cas9 and guide RNA transgenes and Cpf1 – CRISPR RNA ribonucleoprotein complexes, respectively. However, their methods were not described in detail and referred to methods for preparing protoplasts from Arabidopsis leaves or cabbage cotyledons. These recent studies illustrate the value of using protoplasts to rapidly demonstrate the application of new biotechnology tools in soybean. In Arabidopsis, the use of protoplasts has provided important insight into pattern-recognition receptor-triggered and NLR-triggered signaling mechanisms (He *et al*., 2007). Here, we demonstrated that soybean protoplasts are useful for rapidly interrogating the functions of proteins in effector-triggered immune signaling.

Once we confirmed that we could express luciferase in soybean protoplasts, we tested whether *Gm*PBS1-1 or *Gm*PBS1-2 containing a five-alanine insertion at the AvrPphB cleavage site activated cell death, as assessed by a reduction in luciferase expression. These assays showed that both proteins can activate cell death in the absence of AvrPphB (Fig. 2 and Supp. Fig. S4). An equivalent insertion in the Arabidopsis PBS1 protein activates the Arabidopsis RPS5 NLR resistance protein, leading to activation of a hypersensitive response (DeYoung *et al*., 2012; Kim *et al*., 2016); thus, these data strongly suggest that soybean contains a putative NLR protein functionally analogous to RPS5 that is activated by a conformational change in soybean PBS1 proteins. Collectively, these data indicate that it should be possible to engineer novel disease resistance specificities in soybean using a PBS1-based decoy strategy, as was done in Arabidopsis (Kim *et al*., 2016).

To enable recognition of the NIa protease from SMV, we replaced the AvrPphB cleavage site within *Gm*PBS1-1 with a seven-amino acid sequence cleaved by the Nia protease (Fig. 3). Co-expression of this decoy derivative of *Gm*PBS1-1 with the NIa protease triggered cell death in soybean protoplasts, indicating that the PBS1 decoy approach will work in soybean using endogenous *PBS1* genes. Significantly, *Gm*PBS1-2^SMV^ and *Gm*PBS1-3^SMV^ were also cleaved by the NIa protease and can thus likely serve as suitable decoys for the SMV NIa protease (Supp. Fig. S4; Supp. Fig. S5). These data strongly suggest that multiple, synthetic soybean PBS1-based decoys can be deployed in parallel to enable recognition of several soybean pathogens at once. We are now in the process of generating transgenic soybean lines expressing the *Gm*PBS1-1^SMV^ construct.

The evidence presented herein suggests decoy engineering may be an effective strategy to confer resistance against SMV. In support of this expectation, we found that expression of AvrPphB protein from the SMV genome renders SMV avirulent in soybean, whereas a protease inactive AvrPphB mutant does not (Fig. 4). These data thus indicate that the defense responses elicited from AvrPphB-mediated cleavage of the soybean PBS1 proteins is effective against SMV in soybean. Our data are consistent with the observations of Wang *et al*. (2006), who showed that expression of the *P. syringae* AvrB protein from the SMV genome also inhibits systemic SMV infection. Additionally, expression of the *P. syringae* AvrPto protein from a *Potato virus X* (PVX)-based vector elicits defense responses that prevent systemic spread of PVX in tomato (Tobias *et al*., 1999). These data establish that plant disease resistance proteins that normally confer resistance to *P. syringae,* will also confer resistance to viral pathogens when activated, likely due to rapid hypersensitive cell death responses. Such disease resistance proteins are thus ideal targets for engineering broad-spectrum disease resistance to biotrophic pathogens.

PBS1-based decoy engineering may be feasible in diverse crop species beyond soybean. *PBS1* is well conserved among flowering plants, with orthologs present in monocot and dicot crop plant species (Caldwell and Michelmore, 2009). Furthermore, AvrPphB has now been shown to cleave PBS1 orthologous proteins in soybean, barley, and wheat, and to activate immune responses in all three species (Sun *et al*., 2017; Carter *et al*., in press). In barley, this immune response is mediated by an NLR protein designated PBR1 (Carter *et al*., in press). Interestingly, PBR1 appears to have evolved independent from RPS5, thus the ability to recognize PBS1 cleavage has evolved at least twice in flowering plants, suggesting that selection to guard AvrPphB substrates occurs across species. It should thus be possible to introduce novel recognition specificities in most plant species that respond to AvrPphB using synthetic PBS1-based decoys.

## Materials and Methods

### Plant Material and Growth Conditions

*N. benthamiana* seeds were sown in plastic pots containing Pro-Mix B Biofungicide potting mix supplemented with Osmocote slow-release fertilizer (14-14-14) and grown under a 12-h photoperiod at 22°C in growth rooms with average light intensities at plant height of 150 µEinsteins m^-2^ s^-1^

Seed for soybean (*Glycine max* (L.) Merr.) cultivars were ordered from the U.S. Department of Agriculture Soybean Germplasm Collection via the National Plant Germplasm System Web portal (http://www.ars-grin.gov/npgs). Soybean plants were sown in clay pots containing Pro-Mix B Biofungicide potting mix supplemented with Osmocote slow-release fertilizer (14-14-14) and grown in a growth chamber under a 16 hr light/8 hr dark photoperiod at 23°C with average light intensities at plant height of 300 µEinsteins m^-2^ s^-1^.

### *P. syringae* DC3000(D36E) *in planta* Assays

Previously generated plasmids pVSP61-AvrPphB and pVSP61-AvrPphB(C98S) (a protease inactive derivative of AvrPphB) (Simonich and Innes, 1995; Shao *et al*., 2003) were transformed into D36E, a derivative of *Pseudomonas syringae* pv. *tomato* DC3000 with all type III effector genes removed (Wei *et al*., 2015). Bacteria were grown on King’s medium B (KB), supplemented with rifampicin (100 μg/mL) and kanamycin (50 μg/mL), for two days at 30°C. Bacterial lawns of each strain were grown from single colonies selected on KB medium. *P. syringae* DC3000(D36E) strains were resuspended in 10 mM MgCl_2_ to an optical density at 600 nm (OD_600_) of 0.2 for each strain. Bacterial suspensions were infiltrated into the abaxial surface of 14-day old primary leaves of soybean (cv. ‘Flambeau’) seedlings using a 1-mL disposable needleless syringe. Responses were photographed two days after infiltration using a high intensity long-wave (365 nm) ultraviolet lamp (Black-Ray B-100AP, UVP, Upland, CA).

### Phylogenetic Analyses

Soybean PBS1 (*Gm*PBS1) and PBS1-like (*Gm*PBL) homologs were identified by using the SoyBase genome browser (release Williams82.a2.v1; http://soybase.org) (Grant *et al*., 2010) to search the soybean genome with Arabidopsis PBS1 and Arabidopsis PBS1-like proteins (PBL1 through PBL27) as queries. Twenty-two soybean protein sequences were identified as homologous to Arabidopsis PBS1. Amino acid alignments were made using MUSCLE with default parameters. Phylogenetic trees were generated for the collected sequences using the program MEGA7 under a neighbor joining model, and clades were assessed using 1,000 bootstrap repeats (Kumar *et al*., 2016).

### Plasmid Construction and Site-Directed Mutagenesis

The AvrPphB:myc, AvrPphB(C98S):myc, and *At*PBS1:HA constructs have been described previously (Shao *et al*., 2003; Ade *et al*., 2007; DeYoung *et al*., 2012). Glyma.08G360600 (*GmPBS1-1*), Glyma.10G298400 (*GmPBS1-2*), and Glyma.20G249600 (*GmPBS1-3*) were PCR amplified with *attB*-containing primers from soybean cv. ‘Flambeau’ cDNA and then sequenced (see Supplemental Table 1 for list of primers used). These cDNA sequences matched splice variants Glyma.08G360600.3, Glyma.10G298400.1 and Glyma.20G249600.2, respectively, and were also the most similar to Arabidopsis *PBS1* among the splice variants for each gene. The SMV *NIa* protease was PCR amplified from pSMV-34 (Ghabrial *et al*., 1990) using primers designed to introduce *attB* sites. The resulting fragments were gel-purified using the QIAquick gel extraction kit (Qiagen), and recombined into the Gateway entry vector, pBSDONR(P1-P4) using the BP Clonase II kit (Invitrogen) (Qi *et al*., 2012). The resulting constructs were sequence-verified to check for proper sequence and reading frame and subsequently designated pBSDONR(P1-P4):*GmPBS1-1*, pBSDONR(P1-P4):*GmPBS1-2*, pBSDONR(P1-P4):*GmPBS1-3*, and pBSDONR(P1-P4):*NIapro*

To generate the *Gm*PBS1-1^SMV^, *Gm*PBS1-2^SMV^*, Gm*PBS1-3^SMV^*, Gm*PBS1-1^5Ala^, and *Gm*PBS1-2^5Ala^ derivatives, we used an established site-directed mutagenesis PCR protocol using pBSDONR(P1-P4):*GmPBS1-1*, pBSDONR(P1-P4):*GmPBS1-2*, *and* pBSDONR(P1-P4):*GmPBS1-3* as templates (Qi and Scholthof, 2008). The resulting constructs were sequence-verified and designated pBSDONR(P1-P4):*GmPBS1-1^SMV^,* pBSDONR(P1-P4):*GmPBS1-2^SMV^*, pBSDONR(P1-P4):*GmPBS1-3^SMV^*, pBSDONR(P1-P4):*GmPBS1-1^5Ala^,* and pBSDONR(P1-P4):*GmPBS1-2^5Ala^*.

To generate protein fusions with C-terminal epitope tags or fluorescent proteins, the pBSDONR(P1-P4):*GmPBS1-1*, pBSDONR(P1-P4):*GmPBS1-2*, pBSDONR(P1-P4):*GmPBS1-3*, pBSDONR(P1-P4):*GmPBS1-1^SMV^*, pBSDONR(P1-P4):*GmPBS1-2^SMV^*, and pBSDONR(P1-P4):*GmPBS1-3^SMV^* constructs were mixed with either the pBSDONR(P4r-P2):*3xHA* or pBSDONR(P4r-P2):*sYFP* constructs and the Gateway-compatible expression vector pBAV154 [pBAV154 is a derivative of the destination vector pTA7001 and contains a dexamethasone inducible promoter; (Vinatzer *et al*., 2006)] in a 2:2:1 molar ratio. The pBSDONR(P1-P4):*NIapro* construct was mixed with pBSDONR(P4r-P2):*5xmyc* and pBAV154 in a 2:2:1 molar ratio. Plasmids were recombined by the addition of LR Clonase II (Invitrogen) and incubated overnight at 25°C following the manufactures instructions. pBAV154-based DEX-inducible constructs were sequence verified and subsequently used for transient expression assays in *N. benthamiana* (Aoyama and Chua, 1997). The pBSDONR(P4r-P2):*3xHA*, pBSDONR(P4r-P2):*5xmyc*, and pBSDONR(P4r-P2):*sYFP* constructs have been described previously (Qi *et al*., 2012).

The pKEx4tr:*e.v.*, pKEx4tr:*LUC*, and pKEx4tr:*AvrB* constructs have been described previously (Chern *et al*., 1996; Leister *et al*., 1996; Tao *et al*., 2000). To generate the pKEx4tr:*AvrPphB* and pKEx4tr:*AvrPphB(C98S)* constructs*, AvrPphB* and *AvrPphB(C98S)* were PCR-amplified using primers designed to introduce *BamHI* and *NotI* restriction sites at each end and the resulting PCR products were cloned into the *BamHI-NotI* site of pKEx4tr. To generate the pKEx4tr:*GmPBS1-1*, pKEx4tr:*GmPBS1-1^5Ala^*, pKEx4tr:*GmPBS1-1^SMV^*, pKEx4tr:*GmPBS1-2*, pKEx4tr:*GmPBS1-2^5Ala^,* and pKEx4tr:*NIapro* constructs, *GmPBS1-1*, *GmPBS1-1^5Ala^*, *GmPBS1-1^SMV^*, *GmPBS1-2, GmPBS1-2^5Ala^*, and the *NIapro* were PCR amplified using primers designed to introduce *XhoI* and *SacI* restriction sites and cloned into the *XhoI-SacI* site of pKEx4tr. The resulting constructs were sequence-verified to check for proper sequence and reading frame.

The pSMV-Nv::*e.v.* and pSMV-Nv::*GFP* constructs have been described previously (Wang *et al*., 2006). To construct the pSMV-Nv::*AvrPphB* and pSMV-Nv::*AvrPphB(C98S)* clones, *AvrPphB* and *AvrPphB(C98S)* were PCR-amplified using primers designed to introduce an NIa protease recognition site followed by an *AvrII* restriction site (Wang *et al*., 2006). The resulting fragments were gel-purified using the QIAquick gel extraction kit (Qiagen), and subsequently introduced into the *AvrII* restriction site in pSMV-Nv (Fig. 3A). The resulting constructs were sequence-verified to check for proper sequence and reading frame.

### Agrobacterium-mediated Transient Expression Assays in *N. benthamiana*

Transient expression assays were performed as previously described (DeYoung *et al*., 2012; Kim *et al*., 2016). Briefly, the dexamethasone-inducible constructs were mobilized into *A. tumefaciens* strain GV3101(pMP90) and streaked onto Luria-Bertani (LB) agar supplemented with gentamicin sulfate (30μg/mL) and kanamycin (50μg/mL). Single colonies were inoculated into 5 mL of liquid LB containing gentamicin sulfate (30μg/mL) and kanamycin (50μg/mL) and were shaken overnight at 30°C at 250rpm on a New Brunswick rotary shaker. After overnight culture, the bacterial cells were pelleted by centrifuging at 3,000xg for 3 minutes and resuspended in 10 mM MgCl_2_ supplemented with 100 μM acteosyringone (Sigma-Aldrich). The bacterial suspensions were adjusted to an optical density at 600nm (OD_600_) of 0.3 prior to agroinfiltration and incubated for 3 hours at room temperature. For co-expression of multiple constructs, the bacterial suspensions were mixed in equal ratios. Bacterial suspensions were infiltrated by needleless syringe into expanding leaves of 3-week-old *N. benthamiana*. Protein expression was induced 40 hours following agroinfiltration by spraying the leaves with 50 μM dexamethasone supplemented with 0.02% Tween20. Samples were harvested for protein extraction at the indicated time points after dexamethasone application, flash-frozen in liquid nitrogen, and stored at -80°C.

### Immunoblot Analyses of *N. benthamiana* Leaves

For total protein extraction, frozen *N. benthamiana* leaf tissue (0.5g) was ground in two volumes of protein extraction buffer (150 mM NaCl, 50 mM Tris [pH 7.5], 0.1% Nonidet P-40 [Sigma-Aldrich], 1% plant protease inhibitor cocktail [Sigma-Aldrich], and 1% 2,2’-dipyridyl disulfide [Chem-Impex]) using a cold ceramic mortar and pestle. Homogenates were centrifuged at 10,000xg for 10 minutes at 4°C to pellet debris. Eighty microliters of total protein lysate were combined with 20 μL of 5X SDS loading buffer (250 mM Tris-HCl [pH 6.8], 10% SDS (sodium dodecyl sulfate), 30% (v/v) glycerol, 0.05% bromophenol blue and 5% β-mercaptoethanol), and the mixture was boiled at 95°C for 10 minutes. All samples were resolved on a 4-20% gradient Precise^TM^ Protein Gels (Thermo Fisher Scientific, Waltham, MA) and separated at 180 V for 1 hour in 1X Tris/Glycine/SDS running buffer. Total proteins were transferred to a nitrocellulose membrane (GE Water and Process Technologies, Trevose, PA). Ponceau staining was used to confirm equal loading and transfer of protein samples. Membranes were washed with 1X Tris-buffered saline (TBS; 50 mM Tris-HCl, 150 mM NaCl, pH 7.5) solution containing 0.1% Tween 20 (TBST) and blocked with 5% Difco^TM^ Skim Milk (BD, Franklin Lakes, NJ) for one hour at room temperature. Proteins were detected with 1:5,000 diluted peroxidase-conjugated anti-HA antibody (rat monoclonal, Roche, catalog number 12013819001) and a 1:5,000 diluted peroxidase-conjugated anti-c-Myc antibody (mouse monoclonal, Thermo Fisher Scientific, catalog number MA1-81357) for 1 hour and washed three times for 15 minutes in TBST solution. Protein bands were imaged using an Immuno-Star^TM^ Reagents (Bio-Rad, Hercules, CA) and X-ray film.

### Fluorescence Microscopy in *N. benthamiana*

Laser-scanning confocal microscopy assays were performed as previously described (Qi *et al*., 2012). To image protein fusions in live *N. benthamiana* cells, microscopy was performed using an SP5 AOBS inverted confocal microscope (Leica Microsystems) equipped with a 63X numerical aperture 1.2 water objective. The sYFP fusion proteins were excited using a 514nm argon laser and fluorescence detected using a 522-to 545nm band-pass emission filter. mCherry fluorescence (excited with a 561nm helium-neon laser) was detected using a custom 595-to 620nm band-pass emission filter.

### Soybean Protoplast Isolation and Transient Expression Assays

Soybean protoplast isolation and transient expression assays were performed as described previously (Wu and Hanzawa, 2018) with minor modifications. Newly expanded unifoliate leaves from growth chamber (under a 16-h photoperiod at 22°C) grown 12-day-old soybean (cv. Williams 82) were cut into 0.5 – 1 millimeter leaf strips and gently immersed into an enzyme solution [(0.4 M Mannitol, 20 mM MES (pH 5.7), 20 mM KCl, 2% (w/v) Cellulase R-10 (Yakult, catalog number 170221-01, Tokyo, Japan), 0.1% (w/v) Pectolyase Y-23 (Kyowa, catalog number Y-009, Osaka, Japan), 10 mM CaCl_2_, 0.1% (w/v) BSA, 0.5mM DTT) and incubated under vacuum pressure (25 mm Hg) for 30 minutes. Following vacuum infiltration, the leaf strips were incubated in the enzyme solution for 6 hours in the dark at room temperature with gentle agitation (speed = 30, tilt = 1) on a 3-D Rotator Waver (VWR International). After adding 5 mL of W5 solution [(154 mM NaCl, 125 mM CaCl_2_, 2 mM MES (pH 5.7), 5 mM KCl)], the enzyme/protoplast solution was filtered through 75-μm nylon mesh into a 50 mL round bottom tube. The protoplast cells were collected by centrifuging at 100xg for 3 minutes, washed once with W5 solution, and resuspended with MMG solution [(0.4 M Mannitol, 4 mM MES (pH 5.7), 15 mM MgCl_2_)] to the final concentration at 10^6^ mL^-1^ on ice. Five hundred microliters of protoplast cells (5 × 10^5^) were aliquoted and mixed with 50 μg of freshly prepared plasmids and 550 μl of PEG solution [(40% (w/v) PEG4000, 200 mM Mannitol, 100 mM CaCl_2_)] for 15 minutes at room temperature incubation. To stop the transfection, the protoplast cells were washed with 2 mL of W5 solution and resuspended in 500 μl of WI solution [(0.5 M Mannitol, 4 mM MES (pH 5.7), 20 mM KCl)]. After overnight incubation under low fluorescent light (4 µmol m^-2^ s^-1^) and room temperature conditions, the transfected protoplast cells were gently centrifuged (100xg for 3 minutes) and resuspended in 50 μl of WI solution. Ten microliters of substrate solution (ViviRen, E6491, Promega) was mixed with the resuspended protoplasts and the luminescence signal from each sample was recorded using a BioTek Synergy HT plate reader.

### Introduction of SMV-Nv constructs into soybean

The pSMV-Nv::*e.v.*, pSMV-Nv::*GFP*, pSMV-Nv::*AvrPphB*, and pSMV-Nv::*AvrPphB(C98S)* constructs were transformed into *Escherichia coli* TOP10 and streaked onto LB medium supplemented with carbenicillin (100 μg/mL) and 20 mM glucose at 30°C. Single colonies were inoculated into 500 mL of liquid LB supplemented with carbenicillin (100 μg/mL) and 20 mM glucose and shaken overnight at 30°C on a New Brunswick rotary shaker. After overnight culture, plasmid DNAs of pSMV-Nv::*e.v.*, pSMV-Nv::*GFP*, pSMV-Nv::*AvrPphB* and pSMV-Nv::*AvrPphB(C98S)* were prepared using the plasmid Maxiprep Kit (Qiagen).

Introduction of infectious pSMV-Nv and pSMV-Nv-based derivatives into soybean was performed as previously described (Seo *et al*., 2009). Briefly, 10 μg of each infectious cDNA clone was diluted in 50 mM potassium phosphate (pH 7.5) to a total volume of 80 μL and rub-inoculated with carborundum onto the abaxial surface of 14-day old primary leaves of soybean (cv. Flambeau) seedlings. Following mechanical inoculation, plants were maintained in a growth chamber under a 16 hr light/8 hr dark photoperiod at 23°C. Three weeks post-inoculation, the fourth trifoliate leaflet was photographed under white light, flash frozen in liquid nitrogen, and stored at -80°C.

### Immunoblot Analysis of Soybean Leaves

Immunoblot analyses were performed as described previously (Seo *et al*., 2009). For total protein extraction, flash frozen fourth trifoliate leaflets were ground in three volumes of protein extraction buffer (20 mM Tris-HCl [pH 7.5], 300 mM NaCl, 5mM MgCl_2_, 5 mM dithiothreitol, 0.5% Triton X-100, 1% plant protease inhibitor cocktail [Sigma-Aldrich], and 1% 2,2’-dipyridyl disulfide [Chem-Impex]). Homogenates were centrifuged twice at 10,000xg for 10 minutes at 4°C to pellet debris. Total protein concentration was estimated by the Bradford assay (Bradford, 1976). Ten μg of total protein lysate was combined with 5X SDS loading buffer and the mixture was boiled at 95°C for 10 minutes. All samples were resolved on a 4-20% gradient Precise^TM^ Protein Gels (Thermo Fisher Scientific, Waltham, MA) and separated at 185 V for 1 hour in 1X Tris/Glycine/SDS running buffer. Total proteins were transferred to a nitrocellulose membrane (GE Water and Process Technologies, Trevose, PA). Ponceau staining was used to confirm equal loading and transfer of protein samples. Membranes were washed with 1X Tris-buffered saline (TBS; 50 mM Tris-HCl, 150 mM NaCl, pH 7.5) solution containing 0.1% Tween 20 (TBST) and blocked with 5% Difco^TM^ Skim Milk (BD, Franklin Lakes, NJ) overnight at 4°C. Nitrocellulose membranes were incubated with either 1:5,000 monoclonal mouse anti-GFP antibody (Novus Biologicals, catalog number NB600-597, Littleton, CO), 1:5,000 polyclonal rabbit anti-AvrPphB antisera, or 1:10,000 polyclonal rabbit anti-SMV-CP (SMV coat protein) antibody (Hunst and Tolin, 1982) for one hour at room temperature and washed overnight in TBST solution at 4°C. Proteins were detected with either 1:5,000 horseradish peroxidase-conjugated goat anti-mouse antibody (abcam, catalog number ab6789, Cambridge, MA) or 1:5,000 peroxidase-conjugated goat anti-rabbit antibody (abcam, catalog number ab205718, Cambridge, MA) for one hour at room temperature. The nitrocellulose membranes were washed three times for 15 minutes in TBST solution and protein bands were imaged using an Immuno-Star^TM^ Reagents (Bio-Rad, Hercules, CA) or Supersignal^®^ West Femto Maximum Sensitivity Substrates (Thermo Scientific, Waltham, MA) and X-ray film.

## Acknowledgements

The authors thank Alan Eggenberger and John Hill for generously providing the pSMV-Nv and pSMV-Nv::*GFP* plasmids; Said Ghabriel for supplying the pSMV-34 plasmid; Sue Tolin for kindly providing antibody for the SMV coat protein (SMV-CP); Alexandra Margets and Leina Joseph for technical laboratory assistance; the Indiana University Light Microscopy Imaging Center; Morgan Carter and Brody DeYoung for helpful discussions and critical reading of the manuscript; and the U.S. Department of Agriculture Soybean Germplasm Collection for soybean seed. This work was supported by the National Science Foundation Integrative Organismal Systems grant awarded to RWI and SAW (grant no. IOS-1551452) and by funding from BASF Plant Science. MH was supported by a USDA-AFRI predoctoral fellowship.

---

**Supplemental Figure 1.**
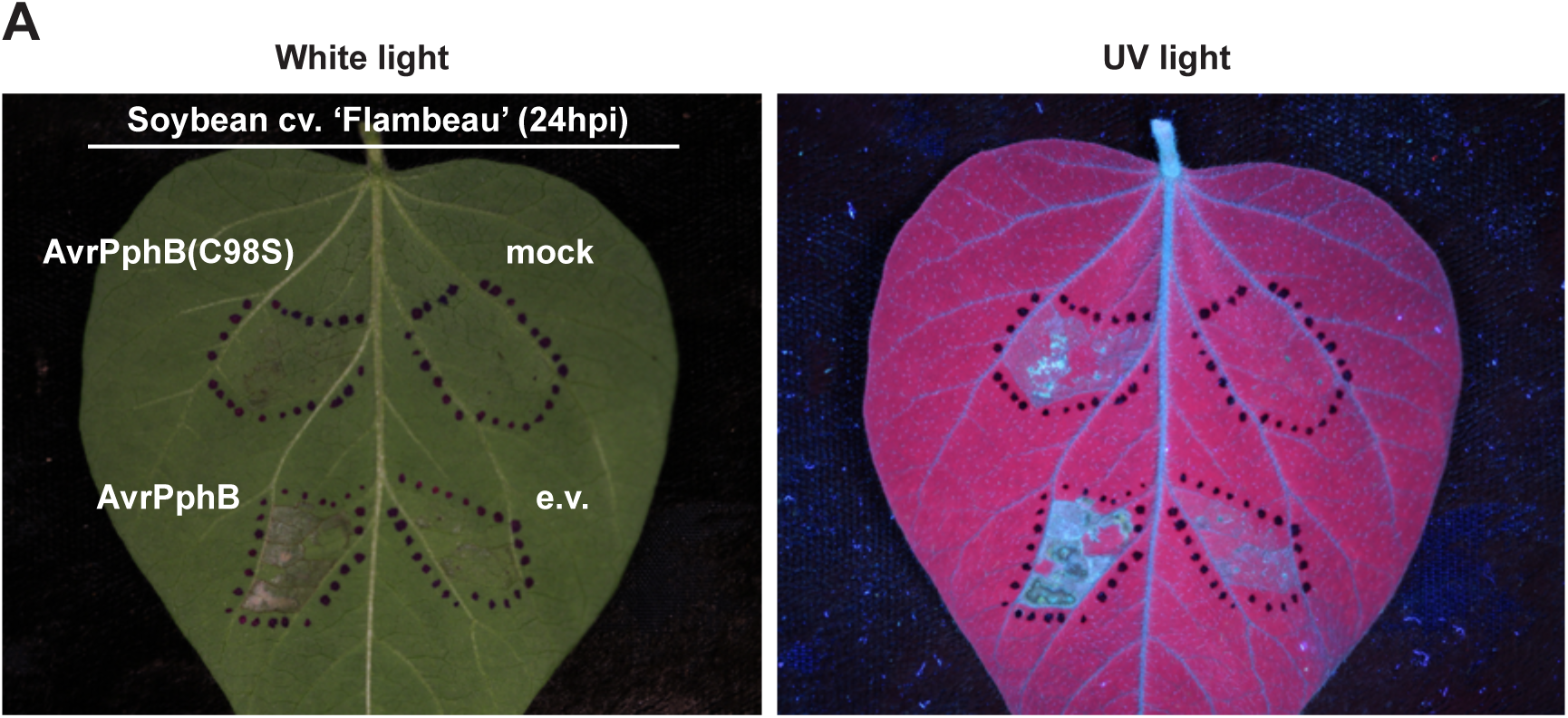
Soybean recognizes AvrPphB protease activity. Response of soybean (cv. Flambeau) to *Pseudomonas syringae* pv. *tomato* DC3000 (D36E) expressing empty vector (e.v.), AvrPphB, or a catalytically inactive mutant of AvrPphB [(AvrPphB(C98S)]. Bacterial suspensions (OD_600_ = 0.2) were infiltrated into the abaxial surface of primary leaves (14-day-old) using a 1-mL disposable syringe. The leaf surface was nicked with a sterile razor blade prior to infiltration. The perimeter of the infiltrated region is indicated with a permanent marker. Photographs were taken twenty-four hours post-inoculation (24hpi) under white light and UV light. A representative leaf is shown. At least five plants were infiltrated with each strain over two repeats.

**Supplemental Figure 2.**
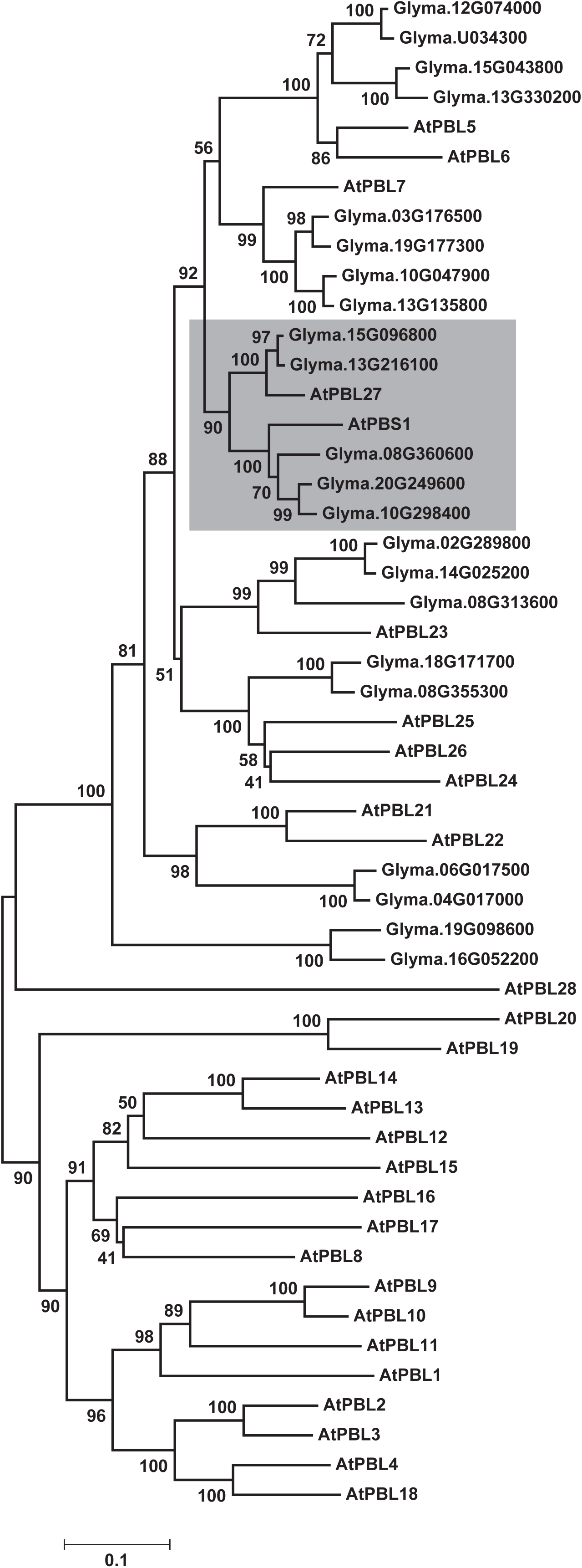
Neighbor-joining phylogenetic tree based on amino acid alignment of full-length products of Arabidopsis *PBS1* (*AtPBS1*), all characterized Arabidopsis *PBS1*-like (*AtPBL*) genes, and soybean *PBS1*-like (*GmPBL*) genes homologous to Arabidopsis *PBS1*. *At*PBS1 and *At*PBL sequences were obtained from The Arabidopsis Information Resource (TAIR10) website (arabidopsis.org; Carter et al., in press). Homology searches were performed using the SoyBase genome browser (release Williams82.a2.v1; http://soybase.org) (Grant et al., 2010) to identify soybean amino acid sequences homologous to Arabidopsis PBS1. Twenty-two soybean protein sequences were identified as homologous to Arabidopsis PBS1. Amino acid alignments were made using MUSCLE with default parameters. The phylogenetic tree was generated for the collected sequences using MEGA7 with the neighbor joining model, and clades were assessed using 1,000 bootstrap repeats (Kumar et al., 2016) (Sun et al., 2017) (Carter et al., in press). The bootstrap values are shown at the nodes. The scale bar indicates amino acid substitutions per site. The gray box highlights the clade presented in Figure 1.

**Supplemental Figure 3.**
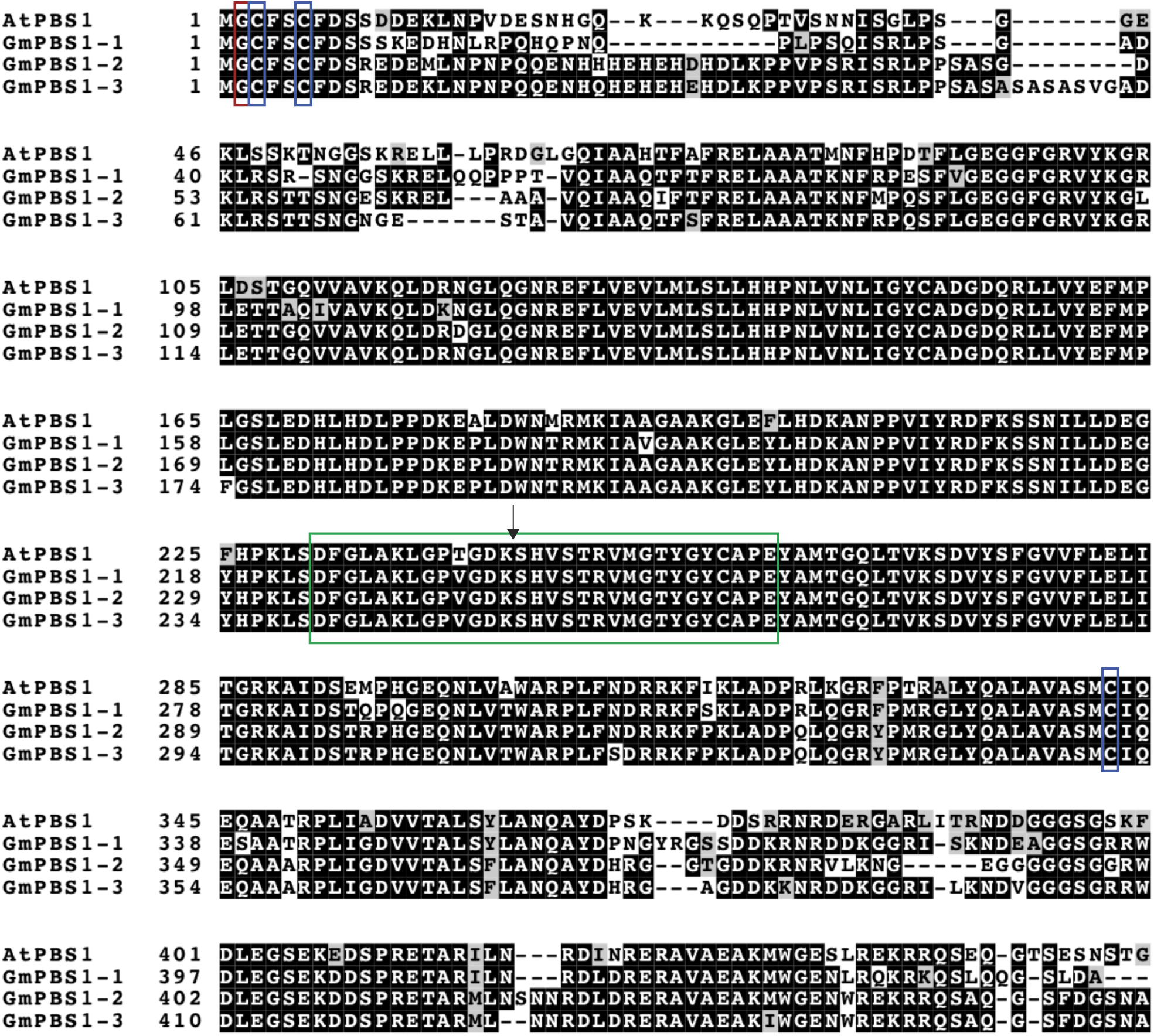
Full-length amino acid sequence alignment between Arabidopsis PBS1 and the soybean PBS1 homologs. Sequence conservation between Arabidopsis PBS1 (*At*PBS1) and the soybean PBS1 orthologous proteins (*Gm*PBS1-1, *Gm*PBS1-2, and *Gm*PBS1-3). Sequence alignment was performed using Clustal Omega (Sievers et al., 2011). Numbers on the left indicate amino acids positions. Conserved amino acid residues and conservative substitutions are shaded in black and grey backgrounds, respectively. Putative myristoylation and palmitoylation sites are indicated with red and blue boxes, respectively. The activation segment is indicated with a green box and the AvrPphB cleavage site with a black arrow.

**Supplemental Figure 4.**
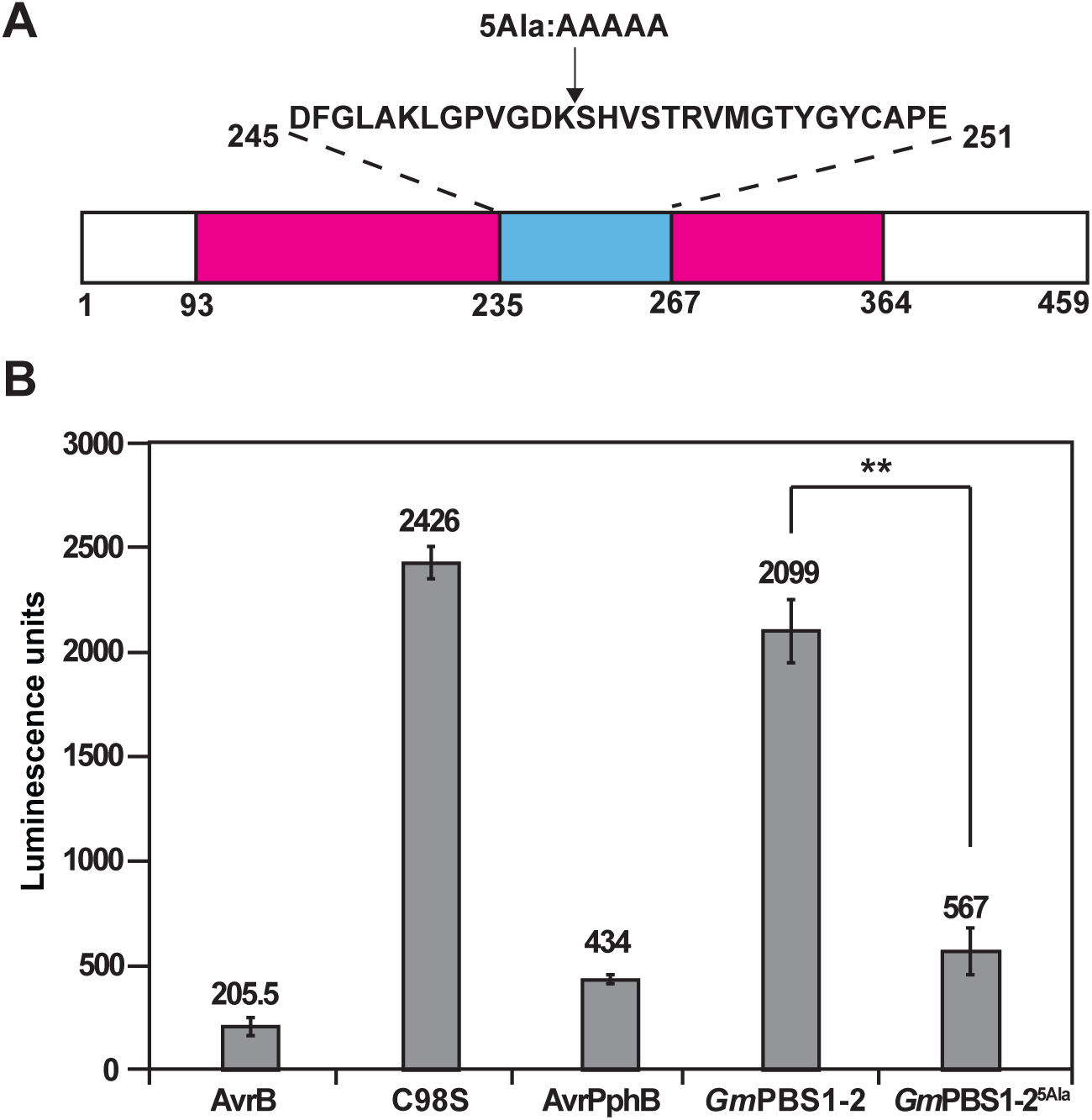
Transient expression of the *Gm*PBS1-2^5Ala^ derivative activates cell death in soybean (cv. Williams 82) A) Schematic representation of the *Gm*PBS1-2^5Ala^ derivative. The predicted kinase domain (amino acids 93-364) and activation segment (amino acids 235-267) of *Gm*PBS1-2 are represented by a magenta box and a cyan blue box, respectively. The amino acid sequence of the activation segment and the location of the five-alanine insertion are indicated above. B) Transient expression of the *Gm*PBS1-2^5Ala^ derivative activates cell death in soybean protoplasts. The indicated constructs were transiently co-expressed along with *Renilla* Luciferase in soybean (cv. Williams 82) protoplasts. Values represent the mean ± S.D. for two technical replicates. T-tests were performed for the pair-wise comparison. The double asterisk indicates significant difference (p < 0.01). Two independent experiments were performed with similar results. The results of one experiment are shown.

**Supplemental Figure 5.**
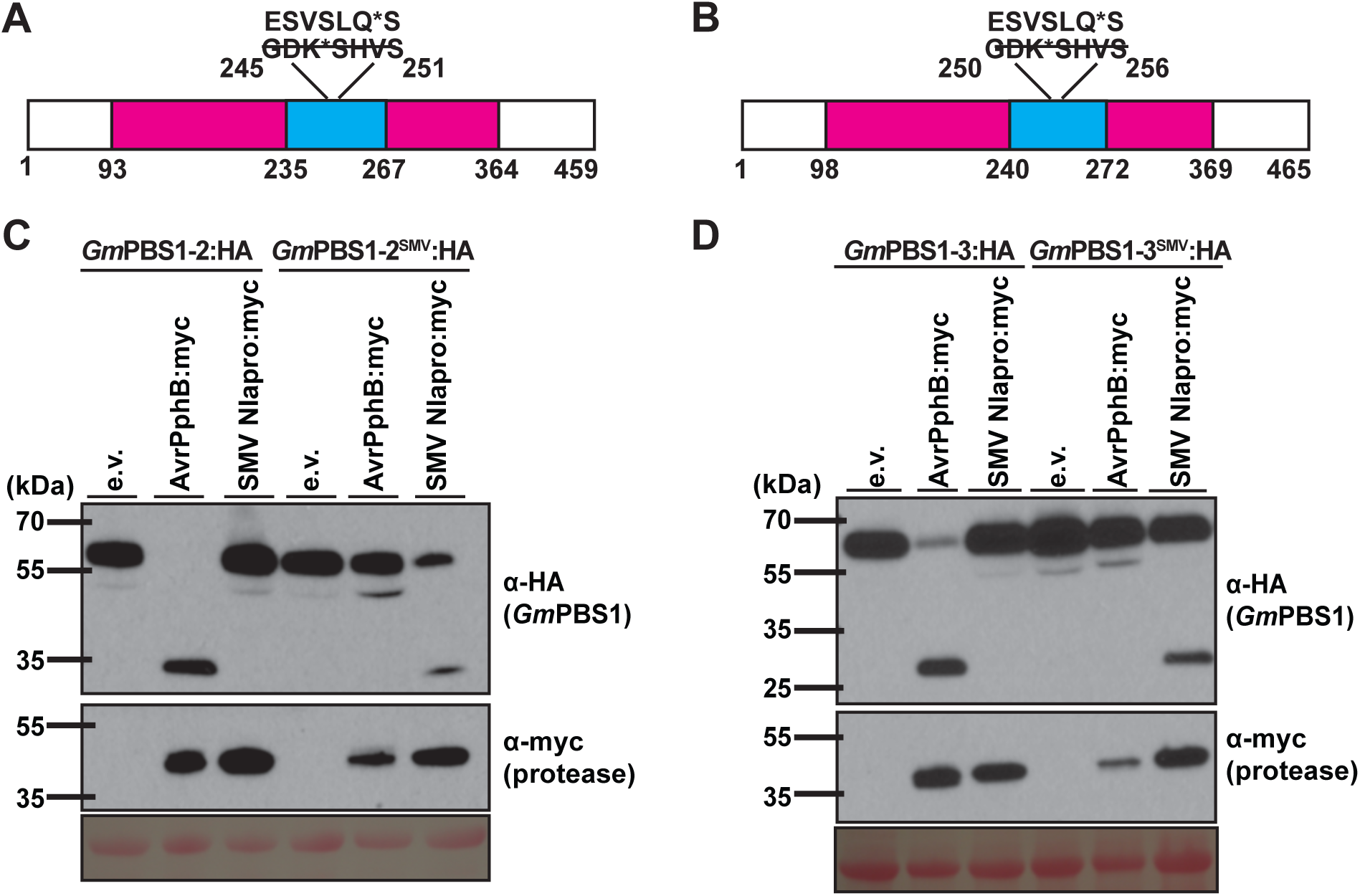
SMV NIa-mediated cleavage of the *Gm*PBS1-2^SMV^ and *Gm*PBS1-3^SMV^ decoy proteins in *N. benthamiana*. A-B) Schematic representation of the synthetic *Gm*PBS1-2^SMV^ and *Gm*PBS1-3^SMV^ decoy proteins. The predicted kinase domain of *Gm*PBS1-2 (amino acids 93-364) and *Gm*PBS1-3 (amino acids 98-369) is represented by a magenta box. The predicted activation segment of *Gm*PBS1-2 (amino acids 235-267) and *Gm*PBS1-3 (amino acids 240-272) is represented by a cyan blue box. The native AvrPphB cleavage site in *Gm*PBS1-2 and *Gm*PBS1-3 (GDKSHVS) was substituted with the cleavage site sequence recognized by the SMV NIa protease (ESVSLQS). The asterisks indicate the location of cleavage by the respective proteases within the recognition sites. C-D) Cleavage of the *Gm*PBS1-2^SMV^ and *Gm*PBS1-3^SMV^ artificial decoy proteins by the SMV NIa protease. HA-tagged *Gm*PBS1-2, *Gm*PBS1-2^SMV^, *Gm*PBS1-3, or *Gm*PBS1-3^SMV^ were transiently co-expressed with either empty vector (e.v.), AvrPphB:myc, or SMV NIapro:myc in *N. benthamiana*. Total protein was extracted nine hours post-transgene induction and immunoblotted with the indicated antibodies. Ponceau S solution staining was included as a control to show equal loading of protein samples. Three independent experiments were performed with similar results. The results of only one experiment are shown.

